# Validation of species-specificity of two commercial antibodies directed against RILP (Rab-interacting lyosomal protein)

**DOI:** 10.1101/2022.11.01.514750

**Authors:** Chan Choo Yap, Laura Digilio, Bettina Winckler

**Author notes:** **Corresponding authors:** Bettina Winckler and Chan Choo Yap, /5526.

## Abstract

RILP is one of many effectors of RAB7 which bind to RAB7 in its activated GTP-bound state. The exact mechanism by which RAB7 effectors interact with RAB7 in time and space is not well understood. One of the known functions of RILP is to recruit dynein to RAB7-positive late endosomes. Dynein has been shown to be responsible for retrograde transport of RAB7-positive late endosomes in neuronal dendrites. We thus became interested in studying RILP in cultured neurons. We herein validate two commonly used anti-RILP antibodies which are commercially available. We find that both recognize only human RILP, but not mouse or rat RILP. These antibodies are thus not suitable for experiments carried out in mouse or rat cells. Furthermore, we find an unexpected difference between neurons and HEK293 cells in their ability to recruit overexpressed RILP to endosomes and cluster them in the cell center.

## Results and Discussion

### Staining with anti-RILP antibodies in neurons is not changed by expression of two shRILP constructs

Two commercially available antibodies against RILP are commonly used in published papers ^5,6^. We obtained both of these from Abcam, ab128616 and ab140188, to determine the localization of endogenous RILP in cultured rat hippocampal neurons. These antibodies are also available from other suppliers (see Materials and Methods). Since RILP is a RAB7 effector ^7,8^, we expected to see staining similar to RAB7 staining, i.e. associated with late endosomes in the soma and the dendrites. When we stained 11 day old (DIV 11) fixed rat hippocampal neurons with ab128616, we observed greatly variable staining, including occasional large punctate staining in the soma as well as staining of structures reminiscent of actin along processes (Fig. 1A). When we carried out the same staining with ab140188, we observed a different staining pattern that appeared more diffuse, and no clear punctate staining was observed (Fig. 1B). In order to determine if any of the observed staining was specific to endogenously expressed RILP, we used two short hairpin plasmids directed against rat RILP to downregulate RILP protein levels. These shRILP constructs have been previously published ^5,9,10^. Expression of shRILP#1-GFP (Fig. 1C,D) or shRILP#2-GFP (Fig. 1E,F) did not change the respective staining patterns of either anti-RILP antibody ab128616 (Fig. 1C,E) or ab140188 (Fig. 1D,F) compared to non-transfected neurons in the same field or to neurons expressing control shCon-GFP (Fig. 1A,B). There are two possible explanations for these observations. Either the antibodies do not recognize rat RILP or endogenous levels of RILP are too low to be detectable by immunofluorescence. We thus tested these two possibilities.

**Figure 1:**
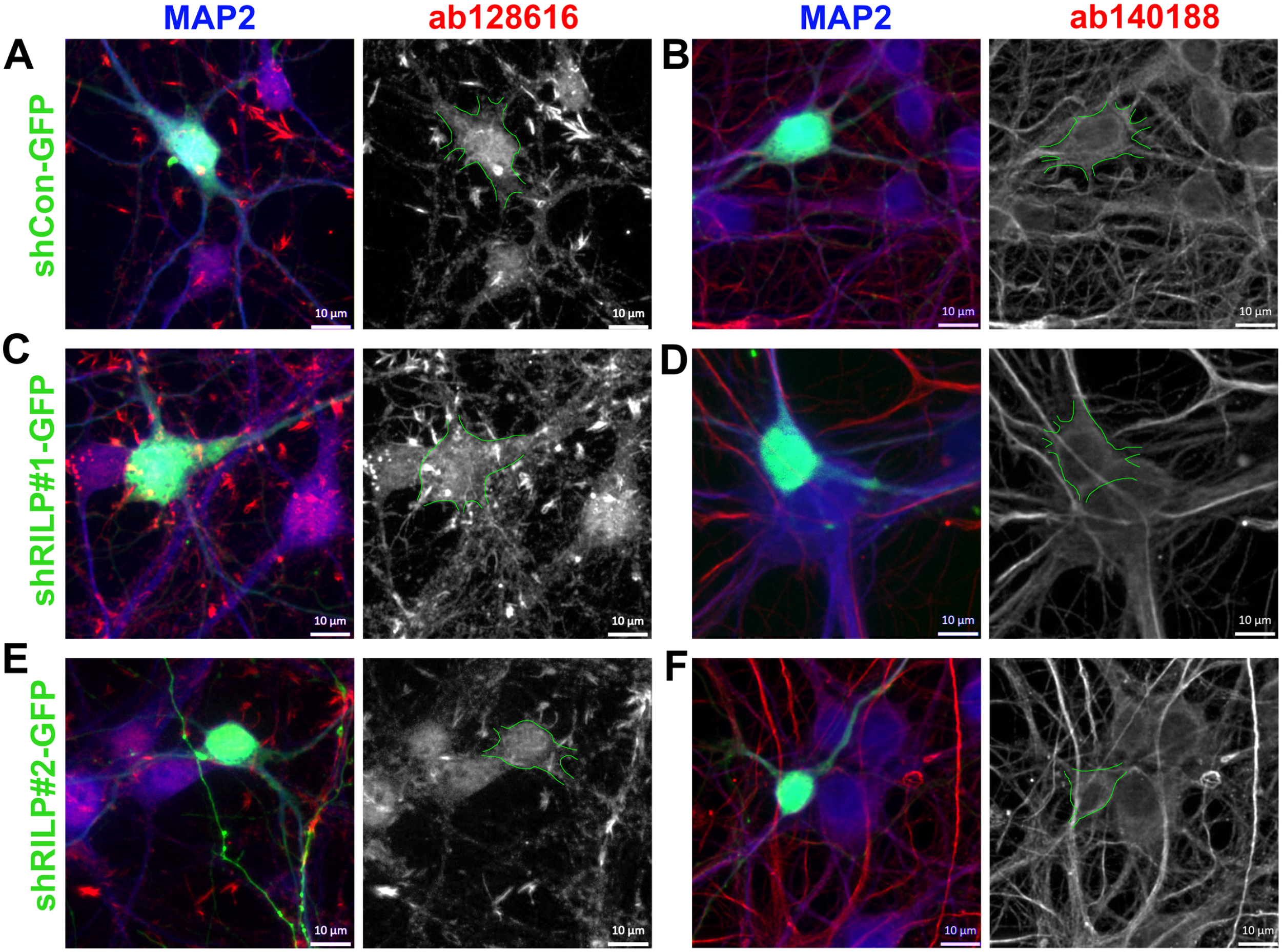
Staining with anti-RILP antibodies in neurons is not changed by expression of two shRILP constructs. DIV5 rat hippocampal neurons were transfected with control shCon-GFP (**A,B**), shRILP#1-GFP (**C,D**) or shRILP#2-GFP (E,F) and fixed 6 days later and stained against MAP2 (blue) to identify dendrites and against RILP using ab128616 (**A,C,E**) or ab140188 (**B,D,F**). The anti-RILP channel is red on the left-hand panel and shown alone as black and white in the right-hand panel. The transfected cell (green) is outlined in green in the black and white panels.

### Commercial anti-RILP antibodies detect clustered human myc-RILP but not overexpressed and clustered rat and mouse RILP

We previously showed that overexpression of human GFP-RILP in rat hippocampal neurons led to perinuclear clustering of late endosomes and lysosomes which were decorated with overexpressed RILP ^4^. We repeated this experiment to test if the commercial anti-RILP antibodies recognized clustered human myc-RILP in cultured neurons. As previously shown for human GFP-RILP, human myc-RILP clustered in the soma (Fig. 2A,B). Counterstaining with ab128616 (Fig. 2A) or ab140188 (Fig. 2B) showed identical clustered staining, indicating that these antibodies recognize human RILP by immunofluorescence.

**Figure 2:**
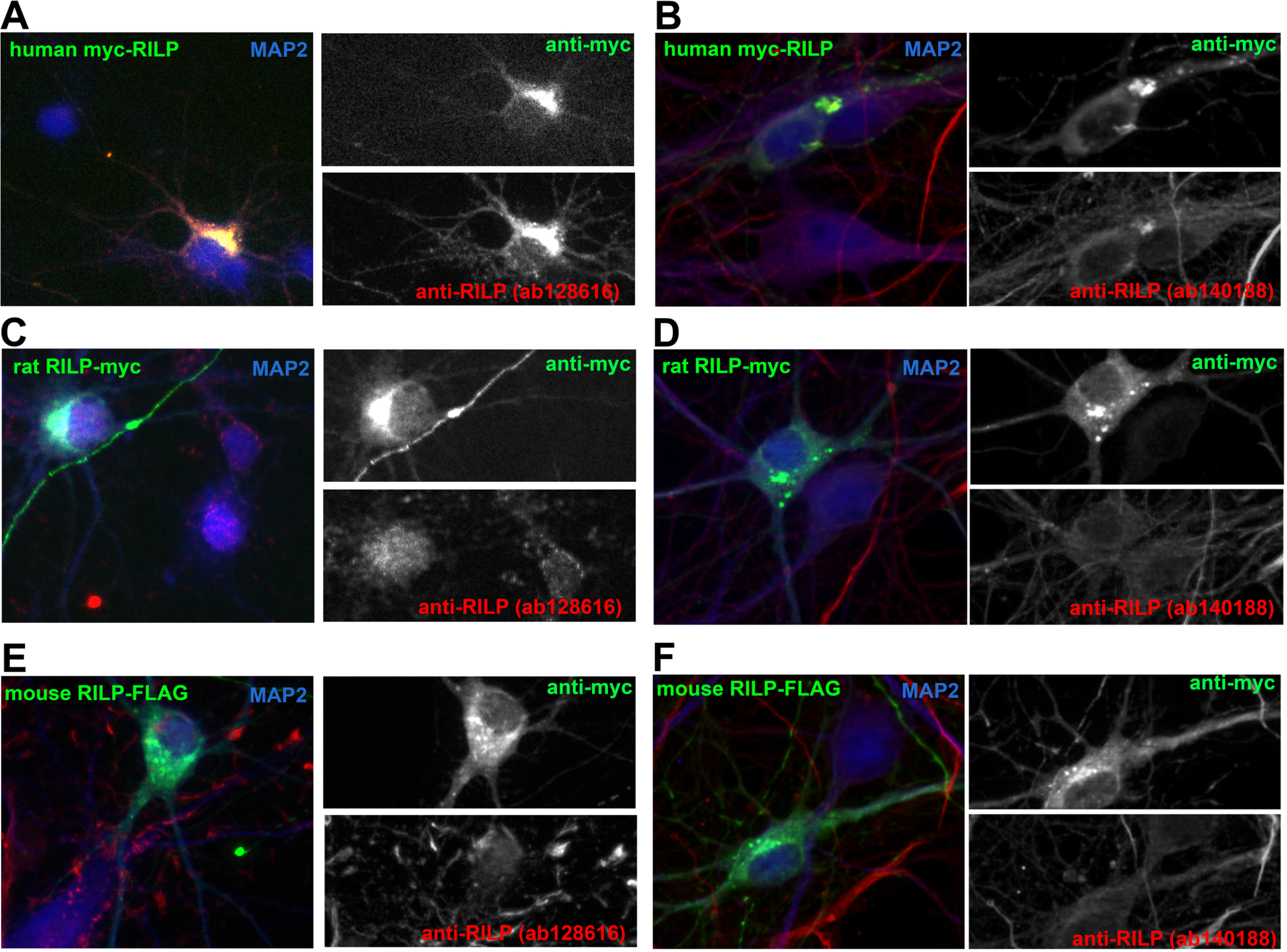
Commercial anti-RILP antibodies detect clustered human myc-RILP but not overexpressed and clustered rat and mouse RILP. DIV7/8 rat hippocampal neurons were transfected with human myc-RILP (**A,B**), rat RILP-myc (**C,D**), or mouse RILP-FLAG (**E,F**). Two days later transfected cultures were fixed and stained against MAP2 (blue) to identify dendrites, against the tag (myc or FLAG; green) and against RILP (red) using either ab128616 (**A,C,E**) or ab140188 (**B,D,F**). Single channel images of the red and green channels are shown in black and white for easier comparison.

Next, we obtained plasmids for rat and mouse RILP, tagged with myc or FLAG at the C-terminus. When we transfected cultured rat neurons with either rat RILP-myc or mouse RILP-FLAG, we again observed striking somatic clustering in the perinuclear region (Fig. 2C-F; green channel). Counterstaining with either ab128616 (Fig. 2C,E) or ab140188 (Fig. 2D,F) showed no clustered staining. In fact, the staining of transfected cells was indistinguishable from untransfected cells on the same coverslip. These observations suggest that these antibodies recognize human RILP by immunofluorescence, but do not recognize rat or mouse RILP.

### Commercial anti-RILP antibodies show only diffuse staining in a human cell line

In order to assess if the two anti-RILP antibodies were able to detect endogenous human RILP by immunofluorescence, we stained the human cell line HEK293. As a positive control, we transfected human myc-RILP. As negative controls, we transfected rat and mouse RILP plasmids, as in Figure 2. Human myc-RILP was often but not always found concentrated in a perinuclear cluster, similar to what has been reported by others ^2,3,11^. Both ab128616 (Fig. 3A) and ab140188 (Fig. 3B) showed higher staining in the transfected cells which co-localized with human myc-RILP (Fig. 3A, green stars). As observed also in cultured neurons (shown in Fig. 2), neither ab128616 nor ab140188 recognized rat RILP-myc (Fig. 3C,D) or mouse RILP-FLAG (Fig. 3E,F). Transfected cells (marked with green stars) were not more brightly stained than neighboring untransfected cells (white outlines). Curiously, neither rat not mouse RILP accumulated in perinuclear clusters when overexpressed in HEK293 cells but instead were found diffusely cytosolic (Fig. 3C-F). This is in contrast with overexpression of any of the tagged RILP constructs (rat, mouse, or human) in neurons (Fig. 2): all accumulated in perinuclear clusters. The molecular basis for this cell type difference is not currently known.

**Figure 3:**
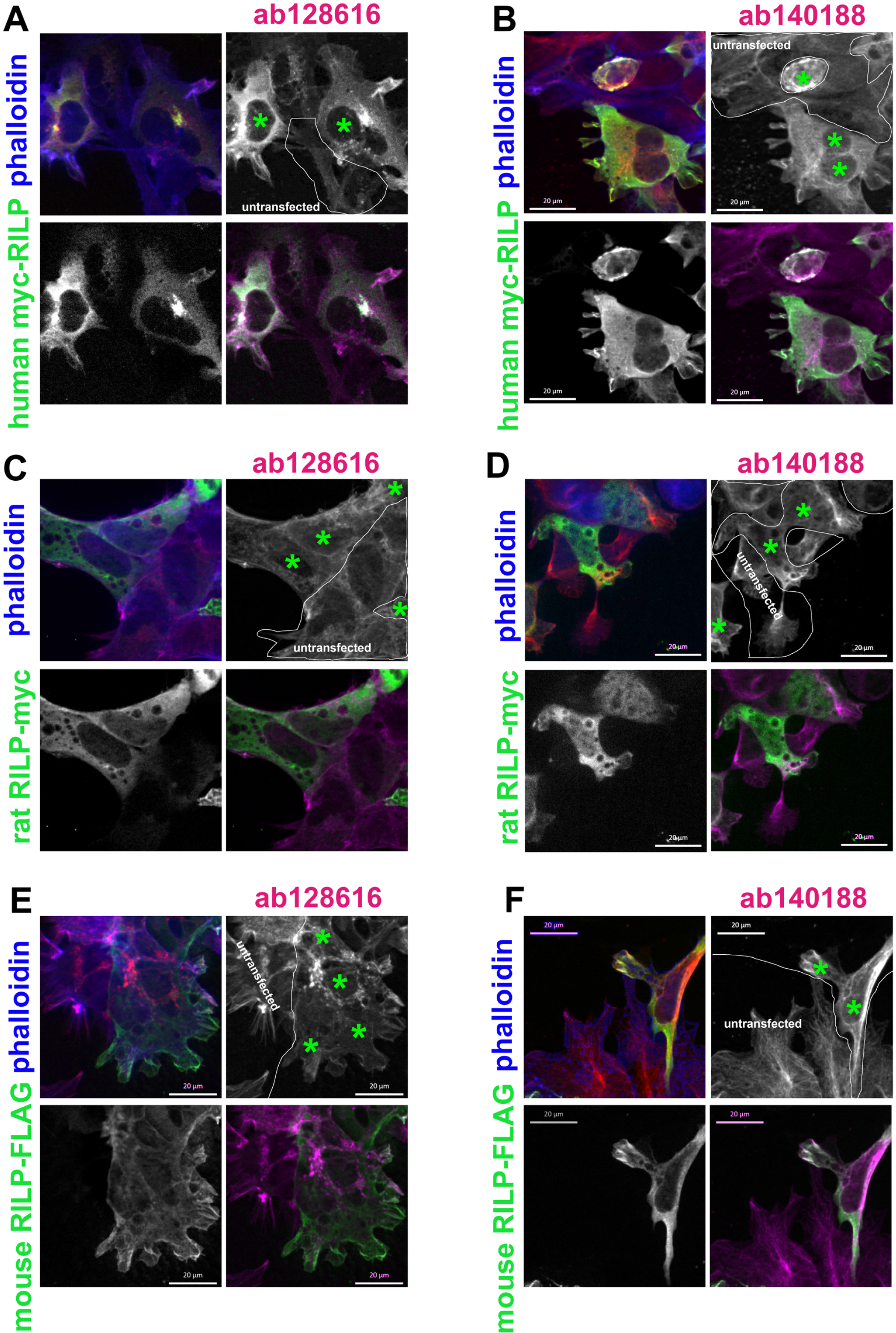
Commercial anti-RILP antibodies show only diffuse staining in a human cell line. HEK293 cells were transfected with human myc-RILP (**A,B**), rat RILP-myc (**C,D**) or mouse RILP-FLAG (E,F) and fixed two days later. Phalloidin (blue) was used to show cell outlines. Fixed cells were co-stained against the tag (myc or FLAG; green)) and against RILP (red) using either ab128616 (**A,C,E**) or ab140188 (**B,D,F**). Single channel images of the red and green channels are shown in black and white for easier comparison. Transfected cells are marked with green stars. Non-transfected cells are outlined in white. The phalloidin channel is omitted in the right bottom corner panels for easier comparison of anti-RILP and anti-tag staining patterns. Only overexpression of human myc-RILP leads to observable clustering near the nucleus in a subset of cells. Neither rat nor mouse RILP overexpression led to clustered distribution. The anti-RILP staining appeared mostly diffuse and no clear endogenous staining pattern could be discerned.

Non-transfected cells showed mostly diffuse staining. Puncta (suggestive of endosomal localization) were rarely, if ever, observed (Fig. 3A,B; white outline demarks non-transfected cells). Given the diffuse staining in HEK293 cells, it is not certain that endogenous expression of RILP is sufficiently high to be detectable. In order to maximize the likelihood to detect endogenous human RILP in HEK293 cells, we overexpressed constitutively active (CA) RAB7 (Figure 4) which leads to high accumulation on late endosomes and recruitment of endogenously expressed RAB7 effectors, including enrichment of RILP. As expected RAB7-CA accumulated on compartments to high levels (Fig. 4A,C and F,H), but staining with ab128616 (Fig. 4A-D) or ab140188 (Fig. 4 F-I) was quite distinct. In order to better compare the perinuclear staining of the anti-RILP antibodies to GFP-RAB7-CA, we created intensity line scans which we forced to run through the brightest RAB7-CA puncta (Fig. 4. E,J). The Rab7-CA peaks (green lines) rarely coincided with high intensity peaks in ab1286161 (Fig. 4E) or ab140188 (Fig. 4J). These observations make it likely that endogenous human RILP is not expressed highly enough to be reliably detectable with these antibodies.

**Figure 4:**
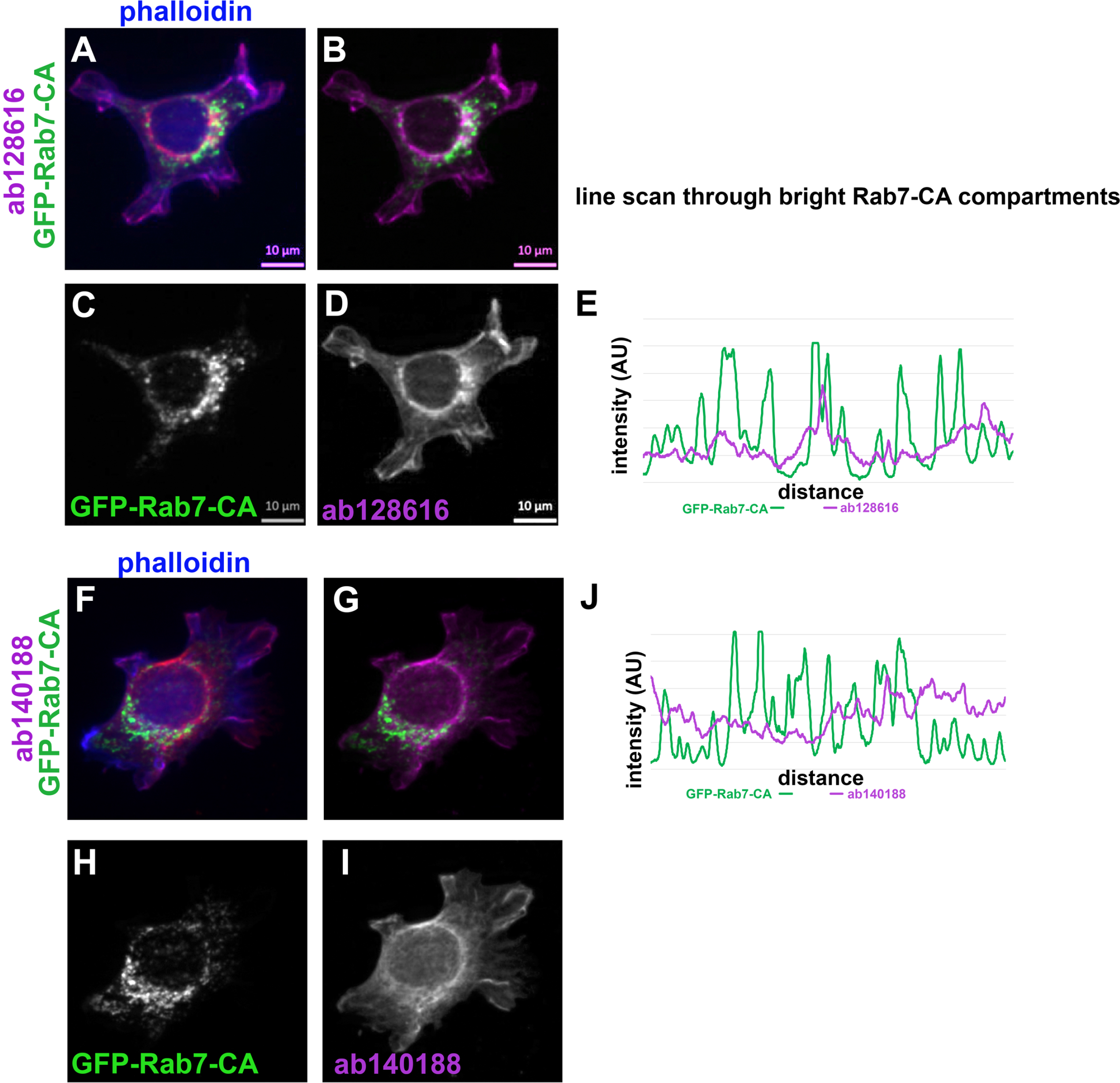
Forced accumulation of activated RAB7 on endosomes does not change staining patterns by ab128616 and ab140188 in HEK293 cells. HEK293 cells were transfected with GFP-RAB7-CA (green; **C,H**) and stained against F-actin (blue, **A,F**) and RILP with either ab128616 (**A-D**) or ab140188 (**F-I**). Single channel images are shown for RAB7-CA (**C,H**) and anti-RILP antibody (**D,I**). A merge of RAB7-CA (green) and anti-RILP (majenta) staining is shown in (**B**) and (**G**). A one pixel wide line was drawn along the bright RAB7-CA compartments to compare the similarity in staining patterns to anti-RILP staining. Intensity profiles along the line are shown in (**E**) for ab128616 and in (**J**) for ab140188.

### Commercial anti-RILP antibodies recognize human but not mouse or rat RILP by Western blot

Lastly, we tested if the commercial anti-RILP antibodies would be able to detect mouse or rat RILP by Western blot. HEK293 cells were transfected with human myc-RILP, rat RILP-myc or mouse RILP-FLAG and lysates were prepared for Western blotting with ab128616 (Fig. 5A) or ab140188 (Fig. 5B). As positive controls, anti-myc or anti-FLAG antibodies were used for Western blotting to detect the transfected proteins in the same lysates (Fig. 5C). ab128616 recognized human myc-RILP (lane 1; Fig. 5A), but not rat or mouse RILP (lanes 3 and 4; Fig. 5A). Non-transfected HEK293 cells (lane 2) showed 2 bands. The expected size of human RILP is around 45kd. The top band is ~75kd and appears to be too large to represent endogenous human RILP. The smaller band appears to be ~40kd and runs substantially below the tagged human RILP. Tt is thus doubtful that it represents endogenous human RILP, but knockdown would be required to conclusively demonstrate this. ab140188 also recognized human myc-RILP (lane 1; Fig. 5B), but not rat or mouse RILP (lanes 3 and 4; Fig. 5B). Non-transfected HEK293 cells (lane 2) showed multiple faint bands and one stronger band which appears to be much too small to be endogenous human RILP. The commercial anti-RILP antibodies thus recognize human RILP by immunofluorescence and Western blotting when overexpressed. RILP is often expressed at very low transcript number, and whether endogenous human RILP is detectable is uncertain. Sequence alignment of human and rat RILP shows that overall sequence homology is only 69% (Fig. 5D). The epitope used to raise ab128616 is known and is underlined in red in Fig. 5D. Fig. 5E shows the alignment of only the 45 amino acid stretch that is the epitope for both human and rat RILP. Sequence identity is only 53%. It is therefore not surprising that ab128616 recognized only human RILP. The exact epitope for ab140188 is not known but corresponds to 18 amino acids in the middle of human RILP.

**Figure 5:**
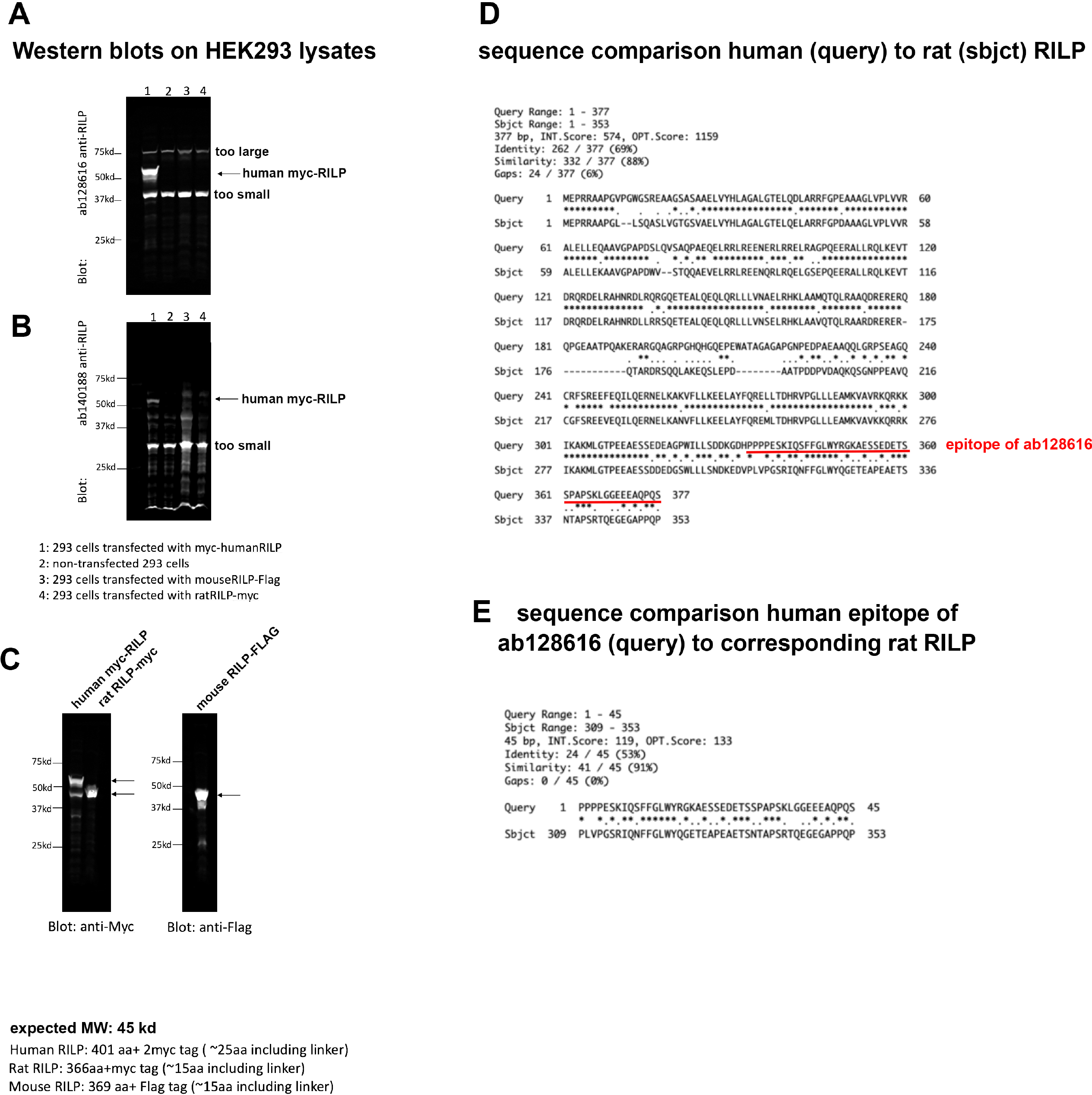
Commercial anti-RILP antibodies recognize human but not mouse or rat RILP by Western blot. (**A,B**) HEK293 cells were transfected with human myc-RILP (lane 1 in A,B), mouse RILP-FLAG (lane 3 in A,B) or rat RILP-myc (lane 4 in A,B). Non-transfected controls were loaded in lane 2. Lysates were probed with ab128616 (A) or ab140188 (B) or against the respective tags (myc or FLAG) in (**C**). Size information is given at the bottom of the figure. Arrows point to the correct bands. Bands of incorrect size are likely not specific. Endogenous RILP is expressed at low levels and likely is not detectable. (**D**)Sequence alignment of human RILP (top sequence) with rat RILP (bottom sequence). The sequence identity is 69%. The epitope of ab128616 is underlined in red. (**E**)Sequence alignment of the epitope used to raise ab128616 comparing human to rat sequence. The identity in this region is 53%.

## Conclusion

In conclusion, we find that two commercial antibodies raised against human RILP only detect human but not mouse or rat RILP protein, both by immunofluorescence and by Western blot. We also find that HEK293 cells have only diffuse staining, and no clear endosomally localized pool of RILP staining can be detected. It is curious that overexpressed human RILP clusters less efficiently in HEK293 cells than in cultured neurons, making it possible that RILP is not efficiently recruited to endosomes in this cell type. Along the same lines, we find that mouse and rat RILP when overexpressed cluster well in the perinuclear region of neurons but are completely diffuse when overexpressed in HEK293 cells. This observation points to neuron-specific mechanisms that enable a high amount of RILP to be recruitable to endosomes. Since RILP is a RAB7 effector, it is possible that higher levels of RAB7 itself contribute to this increased ability of neurons to recruit RILP to endosomes.

## Materials and Methods

### Reagents

| Reagents |  |  |  |
| --- | --- | --- | --- |
| Antibodies | Species | Source, Cat. No | RRID |
| MAP2 | Chicken | EnCor Biotechnology CPCA-MAP2 | AB_2138173 |
| Myc 9E10 | Mouse | Santa Cruz Biotechnology sc-40 | AB_627268 |
| Flag M2 | Mouse | Sigma, F3165 | AB_259529 |
| RILP | Rabbit | Abcam ab128616 (also available from Novus Biologicals # NBP1-98278) | AB_11145272 |
| RILP | Rabbit | Abcam ab140188 (also available from Genetex #GTX32069) |  |
| Alexa 488 anti-mouse | Donkey | Life Technologies A21202 | AB_141607 |
| Alexa 488 anti-rabbit | Donkey | Life Technologies A21206 | AB_2535792 |
| Alexa 568 anti-mouse | Donkey | Life Technologies A10037 | AB_2534013 |
| Alexa 568 anti-rabbit | Donkey | Life Technologies A10042 | AB_2534017 |
| Alexa 647 anti-mouse | Donkey | Life Technologies A31571 | AB_162546 |
| Alexa 647 anti-rabbit | Donkey | Life Technologies A31573 | AB_2536183 |
| Dylight 405 anti-chicken | Donkey | Jackson ImmunoResearch Laboratories 703-475-155 | AB_2340373 |
| Alexa 790 anti-mouse | Donkey | Jackson ImmunoResearch Laboratories 715-655-150 |  |
| Alexa 790 anti-rabbit | Donkey | Jackson ImmunoResearch Laboratories 711-655-162 |  |

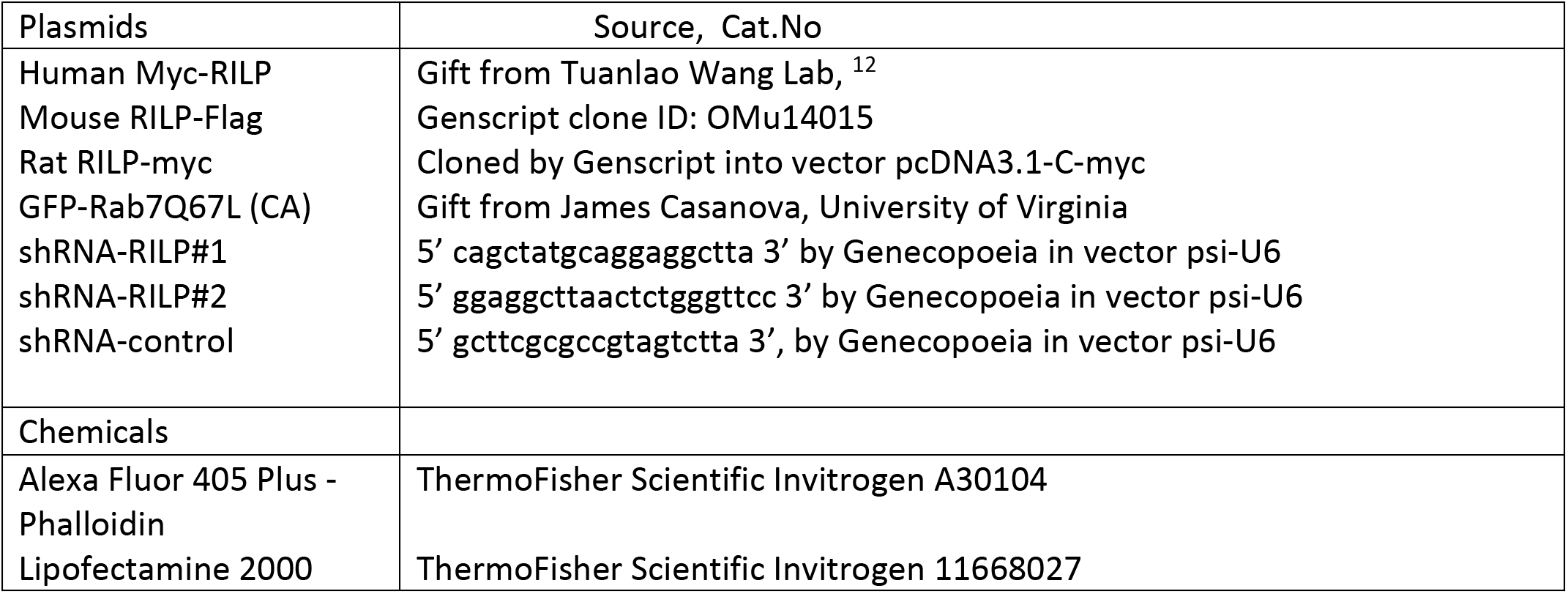

### Neuronal cultures, mammalian cell lines and transfection

Neuronal cultures were prepared as described in ^13^. In brief, the cultures were prepared from E18 rat hippocampi, as approved by the University of Virginia Animal Care and Use Committee. All experiments were performed in accordance with relevant guidelines and regulations (ACUC protocol #3422). Hippocampi from all pups in one litter were combined and thus contained male and female animals. Cells were plated on poly-L-lysine coated coverslips and incubated with plating medium containing DMEM medium with 10% horse serum. After 4 h, the plating medium was removed and replaced with serum-free medium supplemented with B27 (ThermoFisher), and neurons were cultured for 7–10 DIV (days *in vitro*) for experimental use. Transfections were carried out using Lipofectamine 2000 (Invitrogen). Neurons at DIV7-8 were transfected with either myc-HmRILP, RtRILP-myc, MsRILP-Flag, for 36– 40 hours. For RILP knockdown experiments, neurons at DIV5-6 were transfected with shControl, shRILP#1, or shRILP#2 for 6 days. All transfection experiments were repeated in at least 2-3 independently derived cultures.

Hek293 cells were maintained in DMEM+10%FBS. The cells were transfected with human myc-HmRILP, mouse MsRILP-Flag, rat RtRILP-myc, or GFPRab7CA for 48 hours using Lipofectamine2000. All transfection experiments were repeated in at least 2 independently derived cultures.

### Immunocytochemistry

Immunostaining of neurons was carried out as described in ^13,14^. Neurons were fixed in 2% paraformaldehyde/4% sucrose/PBS in 50% conditioned medium at room temperature for 30 minutes, quenched in 10 mM glycine/PBS for 10 minutes. After washing with PBS, cells were then blocked in 5% horse serum/1% BSA/PBS ± 0.2% TritonX-100 for 20 minutes. All antibodies were diluted in 1% BSA/PBS and incubated for 1 hour. Coverslips were mounted in Prolong Gold mounting medium and viewed on a Zeiss Z1-Observer with a 40x objective (EC Plan-Neofluar 40x/0.9 Pol WD = 0.41). Apotome structured illumination was used for most images. Images were captured with the Axiocam 503 camera using Zen software (Zeiss) and processed identically in Adobe Photoshop. No non-linear image adjustments were performed.

### Western Blot

HEK293T cells were transfected with human myc-HmRILP, mouse MsRILP-Flag or rat RtRILP-myc for 3 days and harvested for western blot analysis after lysis in lysis buffer (Cell signaling #9803). Blots were imaged with Li-Cor Odyssey CLx Imager. All western blot analyses were repeated twice.

## Acknowledgments

We thank Dr. Tuanlao Wang (School of Pharmaceutical Sciences, Xiamen University) for providing the human myc-RILP plasmid, and James Casanova (University of Virginia) for GFP-RAB7-CA. This work was supported by NIH R01NS083378 to BW.

